# Combinatorial assembly and design of enzymes

**DOI:** 10.1101/2022.09.17.508230

**Authors:** Rosalie Lipsh-Sokolik, Olga Khersonsky, Sybrin P. Schröder, Casper de Boer, Shlomo-Yakir Hoch, Gideon J. Davies, Hermen S. Overkleeft, Sarel J. Fleishman

## Abstract

Design of structurally diverse enzymes is constrained by long-range interactions that are needed for accurate folding. We introduce an atomistic and machine-learning strategy for Combinatorial Assembly and Design of ENZymes, CADENZ, to design fragments that combine with one another to generate diverse, low-energy structures with stable catalytic constellations. We applied CADENZ to endoxylanases and used activity-based protein profiling to recover thousands of active and structurally diverse enzymes. Functional designs exhibit high active-site preorganization and more stable and compact packing outside the active site. Implementing these lessons into CADENZ led to a tenfold improved hit rate and >10,000 active enzymes. This design-test-learn loop can be applied, in principle, to any modular protein family, yielding huge diversity and general lessons on protein design principles.

## Introduction

Innovation in many areas of engineering relies on the combination of preexisting modular parts(*1*). For instance, in electrical engineering, standard modular parts, such as transistors or processing units, are combined to assemble devices(*2*). Similarly, in a hypothetical, entirely modular protein, fragments could be combined to generate stable, well-folded, and potentially functional domains(*3*). In practice, however, protein domains exhibit a high density of conserved molecular interactions that are necessary for accurate native-state folding. Furthermore, mutations may be epistatic, such that they can only be incorporated against the background of other mutations, severely limiting options for fragment combination(*4*, *5*). Recombination is an important source of novelty in natural and laboratory evolution(*6*–*8*) and the design of *de novo* backbones(*9*); however, due to epistasis, evolution is typically restricted to recombining fragments from only a few high-homology proteins(*6*).

Despite these challenges, immune-system antibodies present a remarkable example in which modularity enables extremely rapid and effective innovation through the combination of a small set of genetic fragments (V, (D), and J genes)(*10*). The result of this process is an enormous diversity of binding proteins that can counter, in principle, any pathogen. Nature has no equivalent strategy to generate structural and functional diversity in enzymes, but some protein folds, such as TIM barrels, β propellers, and repeat proteins, have evolved through the duplication, recombination, and mutation of modular fragments and are therefore prominent candidates for fragment combination. Moreover, these folds comprise some of the most structurally and functionally versatile enzymes and binding proteins in nature(*11*).

Here, we ask whether enzymes could be generated, like antibodies, from combinable fragments? We develop a method, called CADENZ, for Combinatorial Assembly and Design of ENZymes, to design and select protein fragments that, when combined all-against-all, give rise to vast repertoires of low-energy proteins that exhibit high sequence and structural diversity. Isolating active enzymes in such vast protein libraries requires high-throughput screening methods(*12*, *13*), but it can be readily and accurately achieved using activity-based protein profiling (ABPP). ABPP uses mechanism-based, covalent and irreversible inhibitors composed of a chemical scaffold that emulates structural features of the target substrate together with an enzyme active-site electrophile and a fluorophore or affinity tag. To exploit ABPP, we focused on glycoside hydrolase family 10 (GH10) xylanases(*14*–*16*) (Enzyme Classification: 3.2.1.8) as a model system and a dedicated GH10 xylanase-specific activity-based probe (ABP) as the principal enzyme activity readout. We found that CADENZ generated thousands of functional enzymes adopting more than 700 diverse backbones. We then trained a machine-learning model to rank designs based on their structure and energy features. Applying the learned model, we designed a second-generation library that demonstrated an order of magnitude increase in success rate in obtaining functional enzymes.

### Design of modular and combinable protein fragments

For a protein fold to be a candidate for modular assembly and design, its secondary structure elements should be conserved among homologs, but loop regions should exhibit diverse conformations including insertions and deletions(*17*–*19*). In such cases, the secondary structure elements typically provide robustness, whereas the loop regions encode functional differences. The TIM-barrel fold is a prime example of such modularity in which eight β/α segments comprise an inner β barrel surrounded by α helices(*20*, *21*). The catalytic pocket is located at the top of the barrel with critical contributions from all β/α loops. Evolutionary analysis indicates that TIM barrel proteins arose by dual duplication of an ancestral β/α-β/α segment, suggesting that modern TIM barrel enzymes can be segmented into four parts (Fig. 1A)(*22*–*24*). Nevertheless, in the course of evolution, each protein accumulated mutations to adapt the inter-segment interactions for specific functional and stability requirements. Therefore, recombining fragments from existing proteins mostly produces unstable and dysfunctional proteins that require further mutational optimization to turn into stable and active ones(*25*). To address this problem, the CADENZ design objective is to compute a spanning set of backbone fragments that give rise to folded and active proteins when combined all-against-all without requiring further optimization. The primary challenge CADENZ addresses is designing mutually compatible (modular) fragments, *i.e*., ones among which epistasis is minimal.

**Fig. 1.**
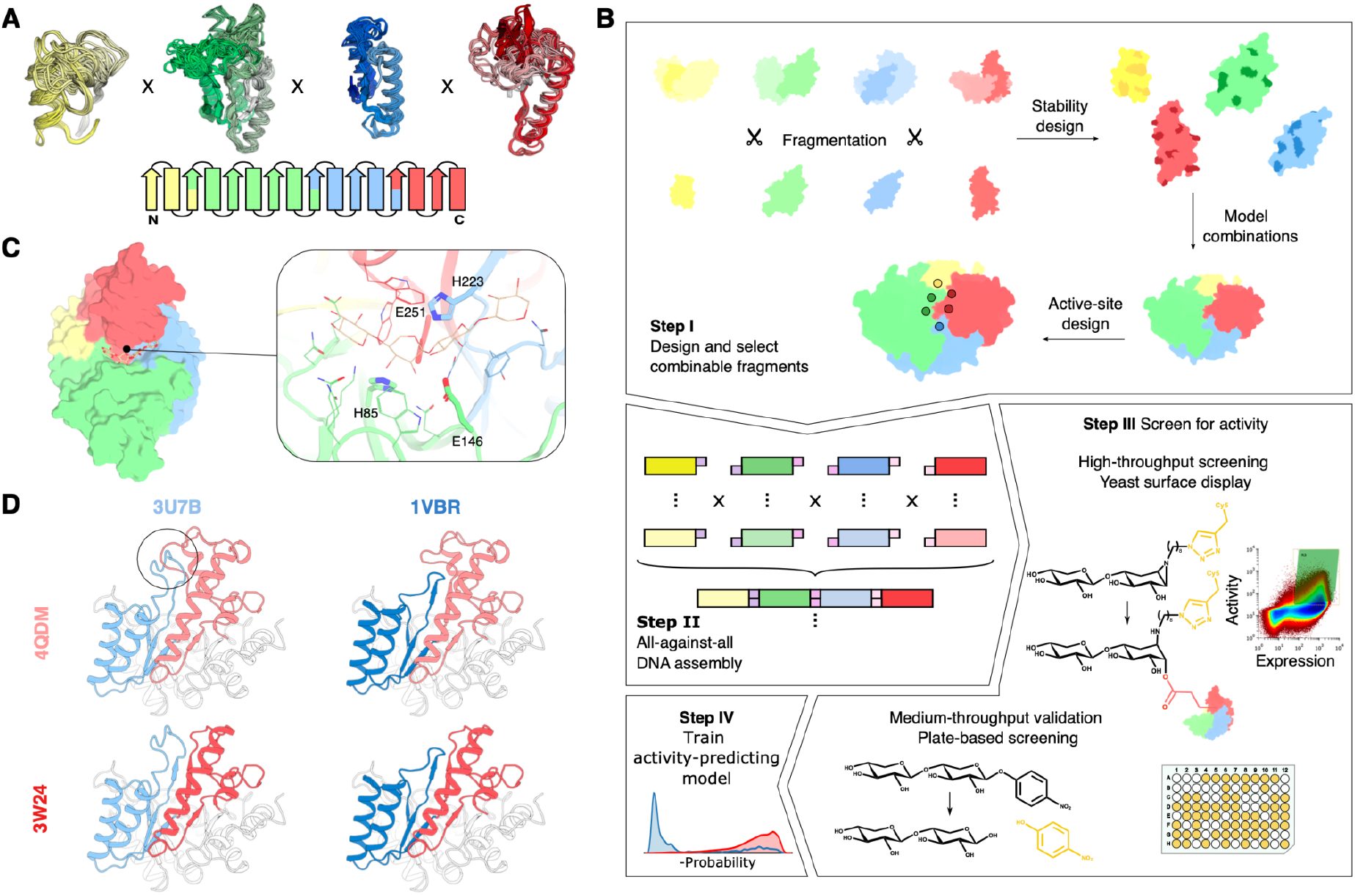
Key steps in the CADENZ workflow. **(A)** (*top*) cartoon representation of selected fragments. (*bottom*) segmentation scheme for GH10 xylanases (color scheme is consistent in all structural figures). **(B)** The design pipeline. **Step I:** Design maximizes internal stability and compatibility with other fragments and diversifies active-site positions that are not directly involved in the catalytic step. **Step II:** DNA oligos encoding fragments are ligated all-against-all using Golden Gate Assembly(*32*) to generate DNA molecules encoding the full-length designs. **Step III:** Designs are sorted using a xylobiose-emulating activity-based probe(*34*) that labels the nucleophilic Glu (red lines) of yeast-displayed functional enzymes. Activity is confirmed on a subset of the selected enzymes in a plate-based chromogenic assay. **Step IV:** An activity predictor is trained based on features that distinguish presumed active and inactive designs. **(C)** Four catalytic residues are restrained throughout design calculations (sticks, numbering corresponds to PDB entry: 3W24). **(D)** Fragments can assemble into low or high-energy structures, depending on other fragments. Segments 3 (blue) and 4 (red) are incompatible (overlap marked by black circle), resulting in extremely high energy (+1,529 Rosetta energy units; R.e.u.) (*top left*). The other designs exhibit low energies (<-950 R.e.u.).

CADENZ starts by aligning homologous but structurally diverse enzymes (in this case, 81 unique structures of GH10 xylanases) and fragmenting them along points that are structurally highly conserved (within the core β segments, see Fig. 1B for a visual guide to the algorithm)(*18*, *26*). Next, the fragments are designed to increase stability while holding the active site fixed. All design calculations take place within a single arbitrarily chosen template (PDB entry 3W24(*27*)) to provide a realistic structural context and promote compatibility between fragments. In practice, each fragment replaces the corresponding one in the template structure, and we use the PROSS stability-design algorithm(*28*) to implement stabilizing mutations within the fragment (8-42 mutations in each fragment; up to 28%) (fig. S1A). In GH10 xylanases, the active site interacts with the xylan substrate through more than a dozen residues from all β/α loops(*29*, *30*) posing a significant challenge for modular design (Fig. 1C). To maintain catalytic activity, in all design calculations the sidechains of four key catalytic amino acids are restrained to their crystallographically observed conformations. At the end of this process, we obtain a set of fragments that are internally stabilized within a common template and designed to support the catalytically competent constellation of active-site residues.

Due to epistasis, however, combining the designed fragments would likely result in mostly high-energy structures that are unlikely to fold into their intended conformation or support the catalytic constellation (Fig. 1D). To address this problem, we enumerate all possible full-length proteins by combining the designed fragments all-against-all and ranking them by Rosetta all-atom energy (Fig. 1B, step I). This process yields hundreds of thousands of unique structures, most of which exhibit unfavorable energies, as expected. To find mutually compatible (modular) fragments, we present a machine-learning-based approach, called EpiNNet (Epistasis Neural Network), which ranks fragments according to their probability to form low-energy full-length structures (Fig. 2A). EpiNNet is trained to predict whether a combination of fragments exhibits favorable Rosetta energy based on its constituent fragments. The trained network weights are then used to nominate fragments to generate the enzyme library. For the following steps, we used the top 6-7 fragments from each segment, assembling into 1,764 structures.

**Fig. 2.**
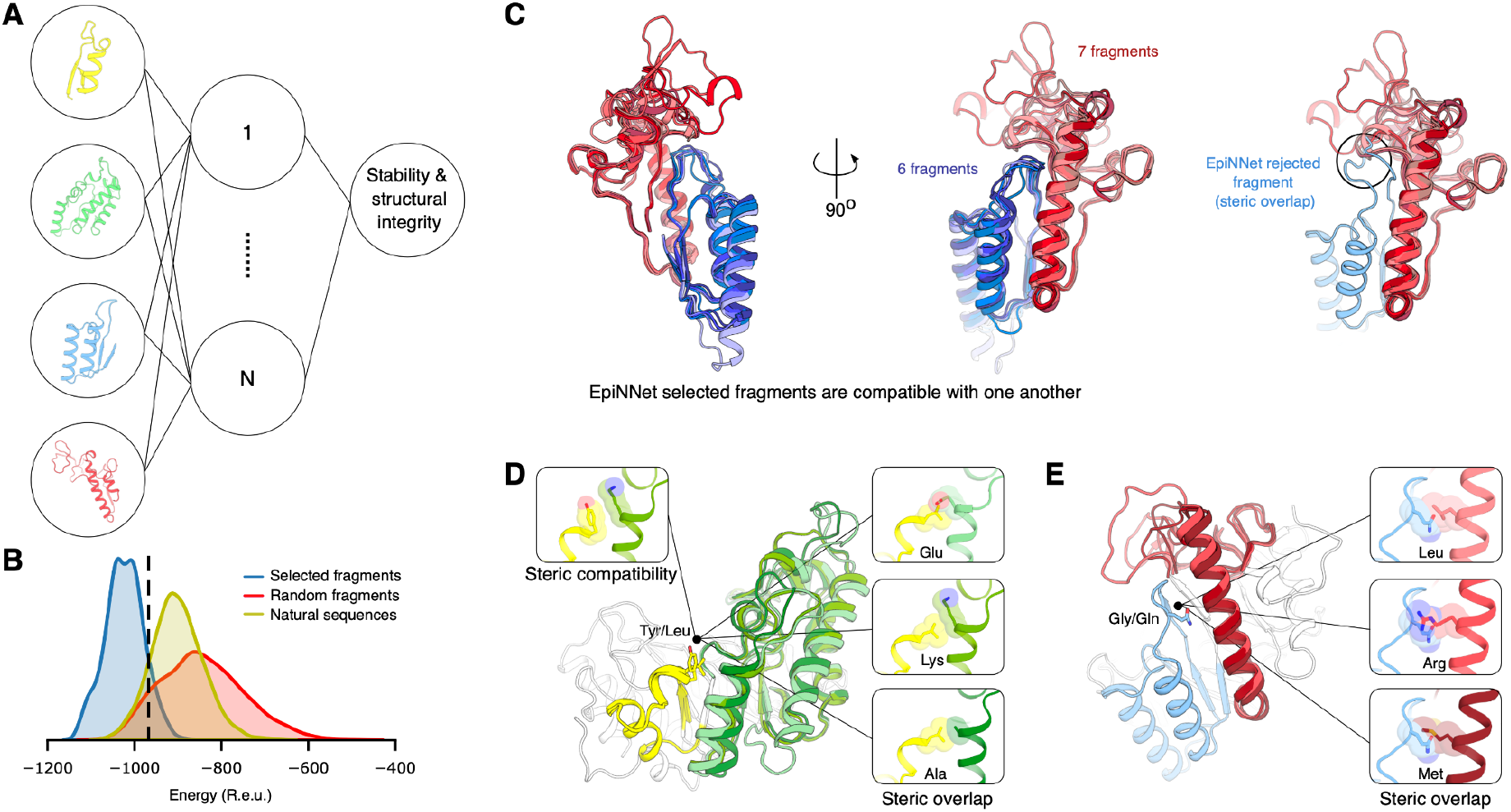
EpiNNet selects fragments that assemble to low-energy structures. **(A)** Schematic representation of the EpiNNet architecture. **(B)** The majority (89%) of the EpiNNet-selected designs exhibit low energy (<-967 R.e.u., dashed line, see Methods) relative to proteins generated by assembling randomly selected fragments or natural ones. **(C)** EpiNNet removes incompatible fragments. (*left*) all fragments selected for segment 3 (blue) and 4 (red). (*right*) discarded fragment with a β/*α* loop that is incompatible with the other fragments. **(D-E)** Examples of mutations selected by EpiNNet (taken from the second-generation library). **(D)** Segment 1 of PDB entry 3W24(*27*) (yellow) faces segment 2 (green); EpiNNet prioritizes a Tyr over Leu which cannot be accommodated with neighboring fragments. **(E)** Segment 3 of PDB entry 1VBR(*55*) (blue) faces segment 4 (red); EpiNNet prioritizes the small Gly over the large Gln.

To add active-site diversity and increase the chances of favorable fragment combination, we next design several sequence variants for each of the backbone fragments. We use the FuncLib design method(*31*) to generate low-energy amino acid constellations at positions in the active-site and in the interfaces between β/α fragments while fixing the conformations of the key catalytic residues as observed in experimentally determined structures (Fig. 1C). We then use EpiNNet again, this time to find the single-point mutations that are most likely to form low-energy full-length proteins in combination with other mutations (fig. S2A).

CADENZ does not necessarily select the lowest-energy fragment combinations but rather mitigates the risk of combining incompatible ones. The consequences of inter-segment epistasis are striking: whereas the energies in the fully enumerated set of designed GH10s can be as high as +2,500 Rosetta energy units (R.e.u.), following EpiNNet fragment selection, the energies are <-890 R.e.u. (Fig. 2B). As a reference, we also generated the distribution of energies obtained by combining the sequence of the fragments selected by EpiNNet prior to any of the design steps. This reference simulates recombination of natural GH10 genes and exhibits a less favorable energy distribution than the combination of PROSS-stabilized fragments (>100 R.e.u. difference on average), underscoring the impact of the design process (Fig. 2B and S1B). Furthermore, EpiNNet alleviates inter-segment epistasis by discarding backbone fragments and designed single-point mutations that are incompatible with neighboring segments (Fig. 2C-E). This analysis highlights the challenge that epistasis poses for effective fragment combination while underscoring the strengths of the EpiNNet selection strategy. Although EpiNNet eliminates more than 60% of the fragments, the designed library exhibits high diversity and includes a total of 952,000 sequences adopting 1,764 different backbones.

### CADENZ generates thousands of structurally diverse and active enzymes

We used Golden Gate Assembly to synthesize full-length genes from the designed fragments(*32*)(Fig. 1B Step II) and transformed the library into yeast cells for functional screening using cell-surface display(*33*) (Fig 1B, step III; see Methods). To probe enzyme activity, we incubated the library with a xylobiose ABP(*34*), which reacts within the enzyme active site to form a covalent and irreversible ester linkage with the glutamic acid nucleophile(*35*). We then used fluorescence-activated cell sorting (FACS) to collect the population of yeast cells expressing active designs (fig. S3A). ABP labeling depends on the nucleophilicity of the catalytic Glu, the ability of the active-site catalytic acid-base residue to enhance the electrophilicity of the ABP-epoxide by protonation, and the integrity of the xylan molecular recognition elements within the active-site pocket. Therefore, this is a sensitive probe for design accuracy in the active site (which comprises elements from all β/α units). Retaining glycosidase ABPs report on the first steps of substrate processing, namely ligand binding to the active site followed by nucleophilic attack. To confirm that selected proteins exhibit the complete catalytic cycle(*36*), we transformed *E. coli* cells with DNA from the sorted population and randomly selected 186 colonies for screening in 96-well plates using the chromogenic substrate 4-nitrophenyl β-xylobioside (O-PNPX_2_)(*37*). This screen demonstrated that 58% processed the substrate (fig. S3B), indicating that most designs selected by the ABP exhibited catalytic activity for this reaction.

We next applied single-molecule real-time (SMRT) long-read sequencing(*38*) to the sorted population. Encouragingly, sequencing showed that the sorted population included a large number of structurally diverse designs: specifically, 3,114 distinct designs based on 756 different backbones (Fig. 3A), compared to only 376 GH10 xylanase entries in the UniProt database(*39*). The recovered designs exhibited many insertions and deletions relative to one another, with sequence lengths varying from 317 to 395 amino acids and 62% sequence identity to one another on average. In all models, residues responsible for the catalytic steps are held in place by construction, but the pocket exhibits high geometric and electrostatic differences (Fig. 3A bottom) due to loop conformation diversity. Strikingly, the designs exhibit as many as 169 mutations and 48-73% sequence identity (Fig. 3B) to their nearest natural homolog (in the nr sequence database(*40*)), and most designs source fragments from four different structures (Fig. 3C).

**Fig. 3.**
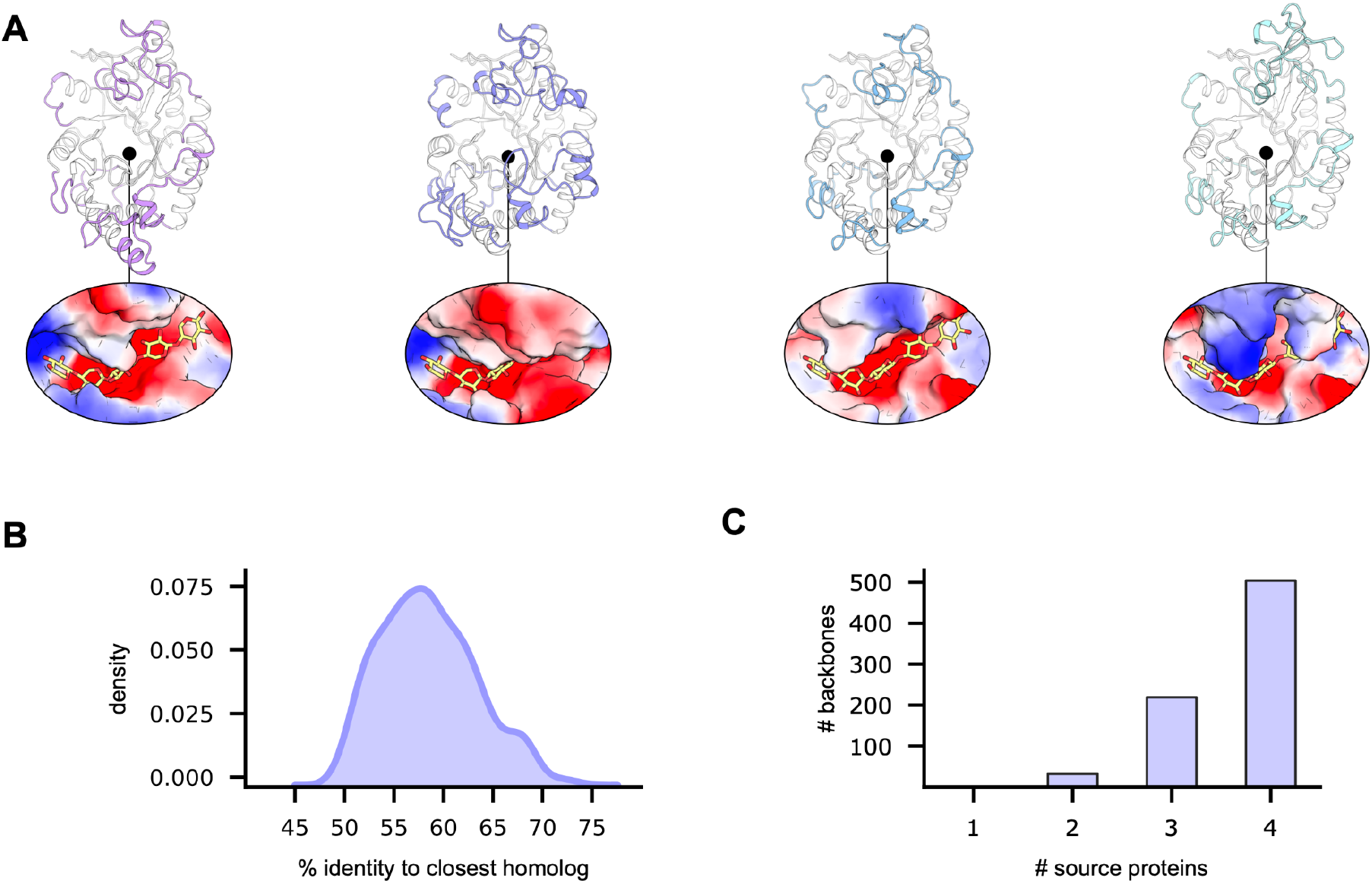
CADENZ generates functional enzymes with high structure and sequence diversity. **(A)** (*top*) representative model structures of recovered enzymes designed by CADENZ. Regions that vary among the four designs are highlighted in colors. (*bottom*) active-site electrostatic potential surfaces of the representative designs exhibit marked differences (putative ligand-bound conformation marked in yellow sticks based on PDB entry 4PUD). **(B)** Distribution of sequence identity to nearest natural homologs of active backbones. **(C)** The number of unique structures from which fragments are sourced. Most active designs incorporate fragments from four different sources.

### Recovered designs are compact and preorganized for activity

The deep sequencing analysis provides a valuable dataset for improving enzyme-design methodology. For each design, we computed 85 structure and energy metrics, some relating to the entire protein, and others restricted to the active site. We avoided using the designed mutations or fragment identities as features for learning so that we might infer general lessons that apply to other enzymes. We tested the differences between the presumed active and inactive sets using an independent two-sample *t*-test, finding that 63 metrics exhibited *p*-values <10^−10^. To select the most meaningful metrics, we visually inspected the distributions of these metrics and focused on ten (Fig. 4A and S4) that do not exhibit significant correlations with one another (Methods). We then trained a logistic regression model based on these ten metrics to predict whether an enzyme is active (Fig. 4B and tables S1–S3).

**Fig. 4.**
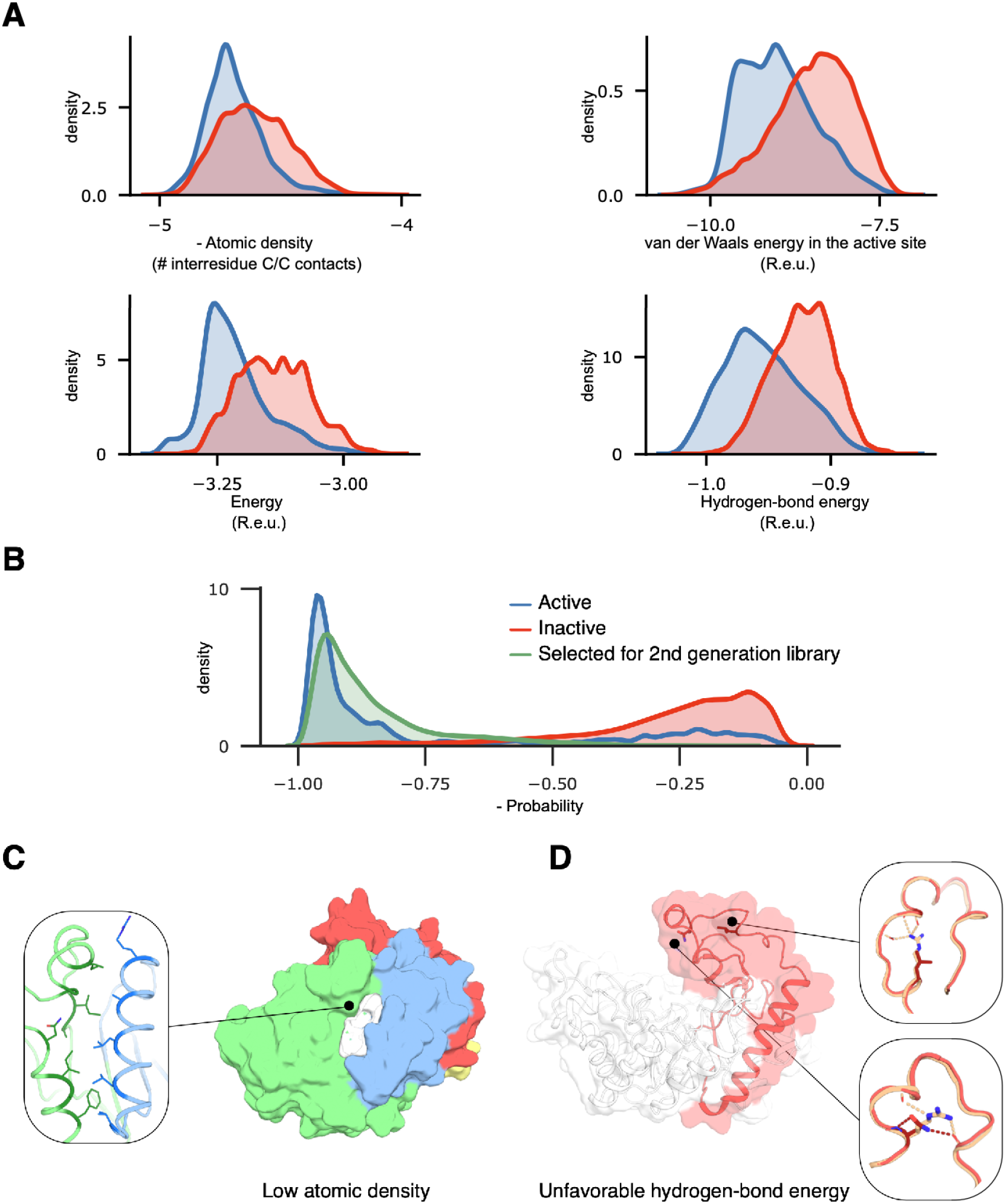
Energy and structure features discriminate between active and inactive designs. **(A)** Representative features included in the activity predictor. The features show a statistically significant difference between presumed active and inactive designs using an independent two-sample *t*-test, but none is individually an effective discriminator. All features are normalized by the protein length. Low values are favorable. **(B)** Separation of designs recovered by ABPP versus other designs based on a logistic regression model. In green, probability distribution for designs assembled by the fragments selected for the second-generation library. **(C-D)** Examples of backbone fragments eliminated by the activity predictor in the second-generation library. Fragment color scheme as in Figure 1. **(C)** 2WYS in segment 2 (green) was selected for the first-generation library but discarded in the second due to low atomic density. The interface with segment 3 (blue) is poorly packed, leaving a gap between the segments (white). **(D)** 1UQZ in segment 4 (red) was selected for the first-generation library but discarded in the second due to unfavorable hydrogen-bond energy. Close inspection revealed two mutations introduced during sequence design, Arg282Asn and Arg289Leu (residue numbering refers to PDB entry: 1UQZ(*56*)), eliminating hydrogen bonds that are crucial for β/α loop backbone stabilization. Mutations in red.

We were encouraged to find that the ten dominant predictive features relate to essential aspects of enzyme catalysis. The most dominant feature is atomic density, which gauges protein compactness and correlates with stable packing. Another dominant feature is the compatibility of the amino acid identity and the local backbone conformation, a key determinant of protein foldability(*41*). By contrast, this feature is disfavored within the active-site pocket, presumably because active-site residues are selected for their impact on activity rather than stability. We also find that hydrogen-bond energies are highly discriminating, reflecting the high propensity of buried long-range hydrogen-bond networks in large proteins of a complex fold such as TIM barrels(*42*, *43*) (fig. S5). Rosetta system energy, however, makes a small contribution to predicting activity, presumably because all designs exhibit low energy by construction (Fig. 2B).

Within the active site, the model assigns almost equal importance to atomic density and van der Waals energy, two features that promote precise catalytic residue placement but penalize overly packed constellations, respectively. The resulting dense yet relaxed packing arrangements are likely to be key to promoting active-site preorganization. Focusing on the four catalytic residues only, the model includes a feature that penalizes high repulsive energy, further underlining the importance of a relaxed and preorganized active site. Our analysis highlights prerequisites of catalytic activity that were not observed in previous high-throughput studies of design methods which focused on the kinetic stability of designed miniproteins and binders(*44*, *45*). We also note that the design objective function is substantially different within the active site versus the remainder of the protein.

Recently, the AlphaFold2 *ab initio* structure prediction method(*46*) has been shown to discriminate correctly from incorrectly folded *de novo* designed binders(*47*). Applying AlphaFold2 to our set, however, showed no discernable difference between presumed active and inactive designs in either the root-mean-square deviation (rmsd) between predicted and designed models or in the AlphaFold2 confidence scores (pLDDT; fig. S6). This result suggests that despite the high mutational load and the sequence and structure diversity in the designs, CADENZ generates sequences with native-like characteristics.

### Order of magnitude increase in design success in second-generation library

We next asked whether the lessons we learned from the first-generation library could improve design success rate. We used the same set of combinatorial designs from the first library, but this time, instead of ranking them based on Rosetta energies, we ranked them according to the activity predictor (Fig. 4B and S2B). Then, we applied EpiNNet to nominate fragments that are likely to be mutually compatible. As in the first library, we designed several sequence variants for each backbone fragment. Here too, sidechain conformations of the core catalytic residues were held fixed in all design calculations. This library included three backbone fragments for each of the four segments and up to 11 sequence variants per fragment (for a total of 100 designed fragments), resulting in 334,125 designed full-length xylanases using 81 different backbones. To gain insight into the molecular features that are disfavored by the activity predictor, we analyzed what backbone fragments were chosen in the first library but discarded in the second. We found, for instance, that atomic density (Fig. 4C) and hydrogen-bond energy (Fig. 4D) were unfavorable in many discarded fragments.

We synthesized and screened the second-generation library as before (fig. S7). Remarkably, sequencing confirmed 9,859 active designs, an order of magnitude increase in the rate of positive hits compared to the first library (Fig. 5A). In addition to the xylobiose screen, we also screened the library using an ABP that is based on cellobiose (Fig. 5B), the disaccharide repeat moiety in cellulose rather than in xylan(*48*). We found 2,778 designs that reacted with the cellobiose ABP but were not sequenced in the library sorted with the xylan ABP, for a total of more than 12,637 active designs (3.8% of the population). To verify that the ABPs selected designs that exhibited the full catalytic cycle, we used plate-based validation with O-PNPX_2_ and cellPNP confirming 85% and 60% of active clones in the xylobiose and cellobiose labeled populations, respectively (fig. S7C,D).

**Fig. 5.**
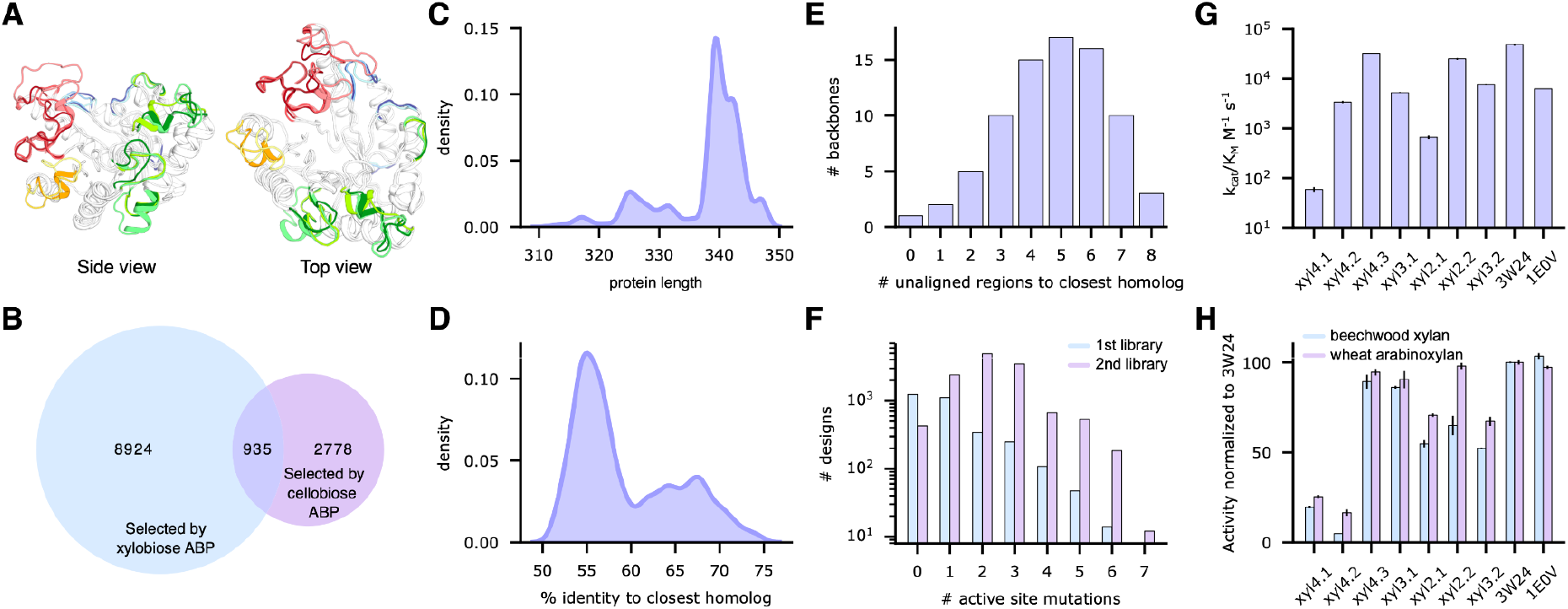
Activity predictor significantly increases design success rate. (**A**) Backbone fragments selected for the second-generation library (colors as in Fig. 1, low diversity regions in white). **(B)** Number of sequences in the population selected by the xylobiose and cellobiose ABPs (blue and purple, respectively). In the overlap region, designs selected by both ABPs. **(C)** Protein sequence length of recovered designs in the second-generation library. **(D)** Distribution of sequence identity to nearest natural homolog of recovered designs in the second-generation library. **(E)** Number of unaligned regions to nearest natural homolog of recovered designs in the second-generation library. **(F)** Designs recovered in the second-generation library incorporate more active-site mutations than in the first-generation library. **(G)** Catalytic efficiency of seven xylanases from the small-set design and two representative natural ones (right, names correspond to PDB entry. 3W24(*27*) is a xylanase from the thermophilic organism *Thermoanaerobacterium saccharolyticum*). The first number in the design name indicates the number of different proteins from which the fragments were sourced. **(H)** Normalized activity with wheat arabinoxylan and beechwood xylan. Data are the means ± standard deviation of duplicate measurements.

Ranking based on the activity predictor resulted in a more focused library that nevertheless includes 79 of the 81 backbones, and sequence lengths ranging from 312 to 347 amino acids (Fig. 5C). Although the activity predictor was blind to the identities of the designed fragments and mutations, we were concerned that it might have focused the second library on a set of fragments identified in functional enzymes in the first library. We analyzed the source of the active designs in the second-generation library, finding that 75% incorporate backbone fragments that were not encoded in the first library and verifying that the learned energy and structure features generalized to fragments not included in the training data. Moreover, the active designs are as divergent from natural GH10 enzymes as in the first library, exhibiting 50-73% identity to the most similar sequences in the nr database (Fig. 5D) and up to 140 mutations and eight unaligned regions (Fig. 5E). Furthermore, the second-generation library incorporates more active-site mutations (Fig. 5F), increasing the potential for altered substrate specificities. We also analyzed the distribution of energy and structure metrics among active and inactive designs in the second-generation library. The discrimination we observed, however, was lower than in the first library, suggesting that the specific learning process we implemented converged.

As an independent test, we applied the learned activity predictor to select a small set of individually designed GH10 enzymes(*18*). Based on the previously described AbDesign strategy(*26*), we used Rosetta atomistic modeling to enumerate all fragment combinations and design their sequences as full-length enzymes, followed by selection using the activity predictor. This strategy encodes more stabilizing inter-segment interactions than when the fragments are designed independently, and the designs are therefore more likely to be stable and foldable. Thus, in this implementation, the design and selection process does not favor modularity but rather optimal structure and energy properties. Twenty-seven designs were selected for experimental characterization with up to 143 mutations and 51-74% sequence identity to their nearest natural homolog. Although these designs were generated using a different process than the one used to train the activity predictor, remarkably, 25 (93%) of the designs were active in hydrolyzing O-PNPX_2_ (table S4), compared to fewer than half in a previous application of AbDesign to GH10 enzymes(*18*). We further characterized the kinetics of the seven most promising designs with various substrates (table S5). Among these designs, several exhibited catalytic efficiencies (*k_cat_/K*_M_) comparable to natural GH10 xylanases from thermophiles, including against natural wood and wheat xylan (Fig. 5G,H) despite incorporating >80 mutations from any known natural protein sequence. These results are a marked improvement in the success of backbone design in enzymes and underscore that the lessons we learned from high-throughput screening can be applied to generate a diverse and highly active set of designs, whether for high- or low-throughput screening.

## Discussion

Modularity is a prerequisite for innovation in numerous engineering disciplines, but protein domains exhibit high epistasis, severely hampering the ability to combine fragments into stable and active structures. CADENZ addresses this conflict by designing a spanning set of low-energy and mutually compatible protein fragments that can be assembled into thousands of diverse and functional proteins. In a companion paper(*49*), we demonstrate that EpiNNet can also be implemented to design large repertoires of active-site sequence variants. This approach therefore increases the number and diversity of functional enzymes that can be interrogated relative to the natural diversity, providing an alternative to metagenomic libraries(*12*). Current methods for optimizing and diversifying proteins rely on sequence statistics(*50*, *51*) or cycles of mutation, recombination, and screening(*52*). Due to high epistasis, these methods explore a small fraction of sequence and structure space, whereas we show in this system that CADENZ can generate 10^6^ structurally diverse designs of which >10,000 are recovered based on activity.

Our results also illustrate that ABPP is an effective strategy for high-throughput isolation of successful CADENZ designs and could be extended to other substrates(*53*), either natural or engineered. The combined strategy enabled us to implement effective design-test-learn cycles on a large number of enzyme designs that have previously led to deeper understanding of the design principles for *de novo* designed miniproteins(*44*, *45*). The rules we learned increased the design success rate by an order of magnitude and were directly transferable to automated small-scale design. Such functional data from many homologous yet structurally diverse enzymes may guide future improvements in macromolecular energy functions and advance efforts to develop AI-based enzyme design methods.

## Acknowledgments

We thank Nir London, Birte Höcker, Shiran Barber-Zucker, and Dina Listov for discussions and Shira Warzsawski and Korina Goldin for technical help. RL-S is supported by a fellowship from the Arianne de Rothschild Women Doctoral Program.

## Funding

Volkswagen Foundation grant 94747 (SJF)

The Israel Science Foundation grant 1844 (SJF)

The European Research Council through a Consolidator Award grant 815379 (SJF)

The Dr. Barry Sherman Institute for Medicinal Chemistry (SJF)

A donation in memory of Sam Switzer (SJF)

The Royal Society for the Ken Murray Research Professorship (GJD)

The European Research Council grant ERC-2011-AdG-290836 ‘Chembiosphing’ (HSO)

The Netherlands Organization for Scientific Research through the NWO TOP grant 2018-714.018.002 ‘Endoglycoprobe’ (HSO)

## Author contributions

Conceptualization: RLS, OK, SJF

Methodology: RLS, OK, SJF

Software: RLS

Validation: RLS, OK

Formal analysis: RLS, SYH

Investigation: RLS, OK, SPS, CDB

Resources: RLS, OK, SPS, CDB, HSO, SJF

Data Curation: RLS

Writing: RLS, SJF, HSO, GJD

Visualization: RLS, OK

Supervision: SJF, HSO

Project administration: SJF

Funding acquisition: SJF, HSO, GJD

## Competing interests

Authors declare that they have no competing interests.

## Data and materials availability

Custom Python scripts, RosettaScripts, commandlines, Jupyter notebooks, and datasets are available at Zenodo(*54*).

## Supplementary Materials

### Materials and Methods

#### Source code

Source code is available at Zenodo(*54*) and in (*26*).

#### Template structure

We selected PDB entry 3W24 as the template structure. Its sequence was designed using PROSS(*28*) using ref2015 as the Rosetta energy function(*57*) with default parameters and ΔΔ*G* cutoff=-0.75 R.e.u. The 3W24 design incorporated 49 mutations. This structure was subsequently idealized and refined (see below). In all following design calculations, the four key catalytic amino acids (residue number: 85, 146, 223 and 251; all numbering with respect to PDB entry 3W24) were restrained to their crystallographically observed side chain conformations. The Rosetta model can be found at <github>/models/3w24_template.pdb.gz

#### GH10 xylanase source structures

We downloaded all GH10 structures from the Pfam database(*58*) (see <github>/data/gh10_pdb_ids) (143 structures). The structures were split into their constituent chains, then idealized and refined using Rosetta (see <github>/rosetta_scripts/idealize_n_refine.xml). A Position-specific Scoring Matrix (PSSM) was generated for each source structure using PROSS(*28*).

#### Source structure segmentation

All source structures were aligned to the template using PyMOL(*59*) and segmented into four fragments. The segmentation points were selected by eye to maximize structural conservation and seamless assembly of different fragments into full length proteins(*26*). The structures were first only crudely segmented, then further refined to match perfectly the template fragments using <github>/scripts/find_best_match.py. The segment borders relative to the template (PDB entry 3W24): S1: 19-45, S2: 46-186, S3: 187-249, S4: 250-320. The PSSMs were segmented accordingly. Example files of the segmented template structure and PSSM can be found at <github>/models/s*_3w24_template.pdb and <github>/pssms/template_s*.pssm.

#### Fragment design

Using AbDesign(*18*, *26*, *43*), each fragment is modeled in the context of the template, and the mainchain dihedral angles are computed and stored for the backbone assembly step (see <github>/rosetta_scripts/splice_out_library.xml for modeling and <github>/models/fragment_design_4pud_s4.pdb.gz for an output example of segment 4 from PDB entry: 4PUD modeled in the context of the template). The segments are two residues shorter than the previous step to allow two constant residues between segments (S1: 20-44, S2: 47-185, S3: 188-248, S4: 251-319). Next, the fragment’s sequence is designed for stability using a custom version of PROSS (<github>/rosetta_scripts/fragment_only_filterscan.xml). In both the modeling and sequence design steps, only the sequence of the fragment is designed, while the template sequence is kept constant.

#### Fragment clustering

To enable full enumeration of all possible fragment combinations and enforce structural diversity, the fragments of each segment were clustered using MaxCluster(*60*) (performing RMSD fit, sequence-independent mode, maximum linkage clustering with the initial threshold set to 0.1 and maximum threshold set to 0.76) resulting in 19-26 clusters per segment. The fragment with the lowest Rosetta score per cluster was selected for the next modeling steps.

#### Modeling all backbone combinations

AbDesign is used to model and refine all backbone combinations (261,326 in total) according to the torsion angles databases and designed sequences from the previous step (<github>/rosetta_scripts/repertoire_splice_in.xml).

#### Select best backbone fragments

The Rosetta scores of the modeling steps are used to train a neural network. In the first library, the designs are labeled “1” if packstat > 0.6(*61*) and Rosetta score is within the top 10% among the designs (< −967.3 R.e.u.). In the second library, we use the whole-structure activity predictor to label “1” designs with predicted probability > 0.88 and packstat >= 0.58 (for labeling with activity predictor: <github>/notebooks/label_using_activity_predictor.ipynb). The network is generated using sklearn MLPClassifier with a single, 4-neuron, hidden layer. We used 4 neurons in the internal-layer to represent the number of segments of the combined structures. The number of neurons can be determined empirically or in relation to the number of combined segments, as we have done here. Post-training, the fragments are ranked according to their activations and the top-ranked ones are selected for sequence design. For GH10, we used 6-7 and 3 fragments per segment for the first and second libraries, respectively. Code, training data and trained model of the first library can be found at <github>/notebooks/select_bb_fragments.ipynb.

#### Computational validation of selected backbone fragment, first library

We compared the energies of the 1,764 selected backbones to: (1) designs generated by randomly selected backbone fragments, where each segment had the same number of fragments as in the first-generation library, including the template’s fragment (for fair comparison). (2) designs generated by the same fragments as in the library, but with the wild type sequence (PROSS sequence for the template). The selected and the random populations had 89% and 9.5% of designs with score better than the “1” labeling cutoff. In the designed backbone with natural fragment sequences, only 2 of the 1,764 designs had better energies than the designed sequence (fig. S1B).

#### Sequence design of selected backbone fragment

For each fragment, we selected 2-11 residues for diversification. We first selected active site residues by adding a xylan substrate from PDB entry 4PUD to all fragments and selecting positions within 3.5 Å from the substrate. For segments 1 and 3 we selected positions on β/*α* loop pointing towards the adjacent segment. We used a custom version of FuncLib(*31*) to calculate the allowed sequence space at the diversified positions for each fragment in the context of the template structure, similarly to the above “fragment design” (<github>/rosetta_scripts/funclib_sequence_space_fragments.xml). In the first library, we used ΔΔ*G*<2.5 R.e.u. and PSSM>0 cutoffs. In the second library there were fewer fragments and thus we considered a larger sequence space: PSSM>-2, ΔΔ*G*<3.5 R.e.u. for segments 1 and 3, and ΔΔ*G*<6 R.e.u. for segments 2 and 4.

To model each fragment’s multipoint mutants in all structural contexts, we model each multipoint mutant on all backbones that can be assembled using the fragments selected in the previous step. Overall, we modeled 1,460,382 and 9,438,696 designs in the first and second library, respectively.

#### Select multipoint mutants

We train a neural network to predict design quality based on the point mutations introduced to the assembled structure. The input layer is a one-hot encoding of the mutations. In the first library, we labeled “1” designs with Rosetta score difference of up to +5 R.e.u. from the no-mutations assembled structures. In the second library, we used the activity predictor with active site features to rank the designs, and labeled “1” the top 10% (*i.e*., probability > 0.63). The network has the same architecture as the network from the best fragment selection step, besides having a hidden layer size matching the number of diversified positions (143 and 65 in the first and second libraries, respectively). Post-training, the multipoint mutants were ranked according to their activations. In the first library, the top-ranked mutants are selected (up to 7 multipoint mutants per fragment). In the second library we observed similar multipoint mutants; thus, to increase diversity, we (1) clustered the multipoint mutants for fragments with >40 multipoint mutants (2) removed backbone fragments without representatives in the top 40% of clustered multipoint mutants, leaving only 3 backbone fragments per segment (3) selected the best (up to 11) multipoint mutants per fragment.

#### Computational validation of selected fragment, first library

To confirm that multipoint mutant combinations give rise to low-energy designs, we randomly modeled 100,000 designs generated by selected multipoint mutants and another set of 100,000 designs from randomly selected multipoint mutants (fig. S2A). While the randomly-selected population had only 62% designs labeled good (defined in the above section), the selected population had 96% good designs.

#### Computational validation of selected fragments, second library

To confirm that combinations of mutated fragments give rise to designs with good predicted probabilities by the activity-predictor, we randomly modeled 50,000 designs generated by selected multipoint mutants and another set of 50,000 designs from randomly selected multipoint mutants (fig. S2B). While the randomly selected population had only 3.9% of designs labeled “1”, the selected population had 95.6% such designs.

#### Design of DNA encoding backbone fragments

We start by designing the DNA to be codon-optimized for *E. coli* expression. To ensure the correct assembly of fragments into full-length enzyme-encoding DNA molecules in the test tube (*i.e*., number and order of fragments), the DNA molecules are flanked with a unique sequence coding for a recognition site for the *BsaI* Type IIs restriction enzyme. This enzyme digests the DNA outside its recognition site, thus leaving a designable 4-bp overhang. DNA molecules encoding for the same segment are designed to have identical overhangs. The DNA sequence of the two-residue constant region between segments is used as overhangs for Golden Gate assembly(*32*). Overhangs are selected to be compatible for one-pot assembly (i.e., overhangs are not reverse complements of one another and at least 2 base pair different from each other). Full DNA sequences can be found at <github>/data/rnd*_fragments_dna.csv

#### Golden Gate Assembly of fragment DNA into full-length enzyme coding molecules

Synthetic dsDNA encoding the fragments were custom synthesized as linear fragments by TWIST Bioscience. All fragments of each segment were pooled. To increase efficiency, we split the Golden gate reaction into three: (1) assembly of fragments from segments 1 & 2 pools; (2) assembly of fragments from segments 3 & 4 pools (3) assembly into final enzyme-encoding DNA by pooling the above two reactions. The assembly is done according to the *Golden Gold (24 fragment) assembly protocol* from NEB(*62*). To avoid non-specific amplification, the assembly product was amplified in two steps: an initial 8-cycles amplification, splitting into 16 reactions, then an additional 8-cycles amplification. Elongation was shortened to 30sec to prevent non-specific amplification.

#### Yeast surface display

The xylanase libraries were cloned into pCTCon2 vector using homologous recombination in yeast, and the active xylanases were selected by yeast-display. The libraries were transformed into EBY100 cells by electroporation(*63*), the cells were grown overnight at 30°C in SDCAA and then resuspended in 10ml induction medium (SGCAA) and induced for 20h at 20°C(*33*). Around 10^7^ cells were taken for labeling, the cells were first labeled with primary antibody for expression monitoring (mouse monoclonal IgG1 anti-cMyc (9E10), Santa Cruz Biotechnology), diluted 1:100 in PBS, for 30 minutes in RT. Then, the cells were labeled with a mixture of secondary antibody for expression monitoring (AlexaFluor488 goat-anti-mouse IgG1 (Life Technologies), diluted 1:100) and 0.03ϻM activity-based probe (xyl-Cy5 or cel-Cy5, diluted from 10mM stock in DMSO), in PBS, for 30 minutes in RT. ABP concentration was chosen by labeling the positive controls (PDB entry: 1TA3 and xyl3.1 and xyl8.3 from(*18*)) with various ABP concentrations, and choosing a concentration at which high signal was obtained while non-specific binding (of a non-expressing population) was minimized. Selections were performed using Fluorescence Activated Cell Sorting (FACS), with BD FACS Melody instrument for the first round library and S3e Cell Sorter (Bio-Rad) for the second round library. Cells were collected based on a combination of three selection gates: yeast cells (FSC vs SSC plot), singlet cells, and 2-5% of top binders. Three sorts were performed for xyl-Cy5 and four sorts for cel-Cy5.

#### O-PNPX_2_ and cellPNP plate-based activity assay

The plasmids from sorted libraries were extracted using Zymoprep Yeast Plasmid Miniprep II kit (Zymo Research). The xylanase inserts were amplified by PCR, re-cloned into pETMBPH vector(*64*) using EcoRI and PstI restriction sites, and transformed into BL21 DE3 cells, which were plated on LB plates with kanamycin. Individual colonies were cultured in 96-well plates with 300 μl of 2YT medium supplemented with kanamycin, the cultures were grown overnight at 37°C and used to inoculate 500μl of 2YT medium supplemented with kanamycin. The cultures were grown for 2 hours at 37°C, induced with 0.2mM IPTG, grown overnight at 20°C, pelleted and frozen at −20°C. The cells were lysed by mixing for 1 hour at 37°C in 300ϻl lysis buffer (Tris pH 7, supplemented with 100 mM NaCl, 0.1mg/ml lysozyme and benzonase) and pelleted. Xylanase activity was assayed using 100μl lysate and 0.5mM O-PNPX_2_ substrate (4-nitrophenyl beta-D-xylobioside, Megazyme) or cellPNP substrate (4-nitrophenyl beta-cellobioside, Sigma Aldrich), at 37°C, by monitoring the absorbance of the leaving group at 405nm in a BioTek plate reader. Designed xylanases xyl3.2, xyl3.1 and xyl3.3(*18*) were used as positive controls. Xylanases were labeled active if the initial reaction velocity (mOD/min) was > 0.3 for O-PNPX_2_ and > 0.4 for cellPNP.

#### Deep sequencing

DNA of the active population was extracted from yeast cells using Zymoprep Yeast Plasmid Miniprep II kit (Zymo Research) and amplified in two steps, as described above for the Golden Gate reaction product. In the second library, each population was amplified with a unique barcoded primer. The *Amplicon Template Preparation and Sequencing* from PacBio(*65*) was used to prepare the DNA for long-read sequencing, and the Circular Consensus Sequence (CCS) analysis with default parameters was used to extract the high-quality reads. Each read was run in BLAST against all possible designs to assign the design it is encoding. In the first library, reads were filtered out if mutated on the protein level relative to their matched design. In the second library, reads were filtered out if #mutations > 2. Reads with #mutations <= 2 were filtered out if mutations were in key positions (i.e., active site and fragment interfaces). The list of active designs can be found here (<github>/data/active_designs_rnd1 and <github>/data/activity_rnd2.pickle).

#### Sequence identity analysis

Sequence identity to closest natural enzymes was calculated using BLASTP(*66*) with the NCBI non-redundant (nr) database (ftp://ftp.ncbi.nlm.nih.gov/blast/db/).

#### AlphaFold2 analysis of designed backbones

The structures of the backbones (without multipoint mutants) in the first designed library were predicted using ColabFold(*67*) with default parameters. RMSD between Rosetta models and AlphaFold2 predictions were done using the RMSD filter in RosettaScripts(*61*).

#### Metric calculations and filtering

All designs in the library were modeled in Rosetta (using <github>/rosetta_scripts/funclib_sequence_space_fragments.xml) and the different metrics were calculated (see <github>/rosetta_scripts/features.xml and <github>/rosetta_scripts/features_repack_activesite.xml for calculations, <github>/data/rnd*_all_seqs.fa for sequences and <github>/data/rnd*_features.csv for metrics data on all designs of the first and second library). All metrics depending on sequence length were normalized (*i.e*., metric / #residues); features relating to the entire protein were normalized by the protein’s length, whereas features restricted to the active site were normalized by the number of residues included in the active site (12-18 residues, depending on the backbone fragments included in the protein). Following visual inspection of the distribution of the metrics in the active and inactive populations, 18 metrics were considered as features for the activity predictor: fa_rep_catres, vdw, vdw_activesite, fa_sol_catres, fa_elec, fa_elec_activesite, hbond, hbond_activesite, p_aa_pp, p_aa_pp_activesite, p_aa_pp_catres, fa_dun_catres, yhh_planarity, yhh_planarity_activesite, degree_activesite, contacts, contacts_activesite, and total_score. The VDW metric is defined as fa_atr + 0.55 * fa_rep (as weighted in Rosetta ref15 scoring function(*57*). The hbond metric is defined as the sum of all hydrogen bond energies (hbond_lr_bb + hbond_sr_bb + hbond_bb_sc + hbond_sc). The features were standardized by removing the mean and scaling to unit variance.

To eliminate correlated features, we used feature selection procedures. The data were split into training and test sets (80 and 20 percent of the data, respectively). We used stepwise selection: at each iteration, there is a forward and a backward step. Forward step: add the feature having the lowest *p*-value when added to the logistic regression model, stop when *p*-value > 0.05 or the feature’s coefficient < 0.1. Backward step: train the model with the newly added feature, and drop features with an increased *p*-values (> 0.05). The stepwise selection procedure was repeated five times, each with a different under-sampled population from the train set. We used the 10 features that were selected in all five repeats. To make sure all features contribute to the model performance, we performed a likelihood ratio test for each feature where we compare the model to a restricted model lacking the tested feature (*i.e*., restricted model). We compute 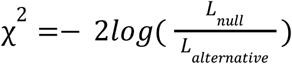. The null hypothesis, where the restricted model is better, is rejected if χ^2^ is greater than a χ^2^ percentile with *k* degrees of freedom (*k* is the difference in parameters between the models; in this case *k*=1). We trained the models using the python package statsmodels, which also computes the log-likelihood of the model. All features significantly rejected the null hypothesis (*p*-value → 0). The features included in the models and their coefficients are summarized in Table S1.

#### Logistic regression model training

A few resampling procedures were tested as the data are highly imbalanced: oversampling, undersampling, and Synthetic Minority Over-sampling Technique (SMOTE), with and without Tomek links (all were performed using the python package imbalanced-learn). All methods gave similar results; thus, we chose random undersampling as it is less prone to overfitting and is faster to train on. We trained two logistic regression models; one including the 10 selected features described above (Logit model from python’s statsmodels, regularized fit with *α*=0.1). The second model includes only features describing all residues of the protein. The features of the second model were selected as described above and are listed in Table S1. The trained models can be found at <github>/ml_models/logit_xylanases*.pickle

#### Design of a small set of xylanases

Designs were generated as described(*26*). Briefly, we used the same structures and PSSMs from the library design. Fragments were extracted and modeled in the context of the template to compute the mainchain dihedral angles using AbDesign (<github>/rosetta_scripts/small_set_splice_out.xml). The fragments of each segment were clustered, and the lowest Rosetta energy fragment was selected for the backbone recombination and design step (25, 20, 17, and 21 fragments for segments 1,2,3, and 4, respectively, see <github>/rosetta_scripts/small_set_splice_in.xml). During design, the active site residues (both core catalytic and binding substrate residues) were biased towards their crystallographically observed conformations. The assembled and designed structures were filtered for: (1) correct positioning of Trp292 and Trp300 in the active site, i.e., 1.5Å RMSD from the conformation in the template structure (2) packstat >= 0.58. The filtered 29,451 designs (18.5% of designed chimeras) were labeled using both activity predictors and 1170 designs ranked among the top 7% of both predictors were selected for PROSS stability design. To control the number of different source proteins in the chimeras, 21 and 72 designs from only 2 or 3 sources (template and either segment 2 or 4 from a different source) were also selected for PROSS. Segments 2 and 4 were selected for diversification as they harbor most active site residues and are more structurally diverse than segments 1 and 3.

We applied PROSS to design the selected designs using a sequence space of mutations with PSSM probability>0 and ΔΔ*G* < −0.75 R.e.u., incorporating 4.6 mutations on average. We filtered the designs again using packstat and Trp residues positioning as described above, and scored the remaining 1,155 designs using both activity predictors. To increase diversity in the experimentally tested set, the designs were structurally clustered (see Fragments clustering above), and a representative with the highest average probability of both activity predictors was selected. We filtered out: (1) representatives with predicted probabilities lower than the mean probability (2) representatives with segments 2 and 4 sourced from the same proteins. We similarly selected designs for the sets of designs where we explicitly select for only 2 and 3-source, filtering out for probabilities lower than the 25th percentile instead of the 50th. Overall, we selected 27 designs; 19, 3 and 5 from general selection, 2-source and 3-source designs, respectively.

We used pSUFER(*68*) to detect poorly designed positions, flagging positions with more than four alternative amino acids with ΔΔ*G* < 0, located outside the active site. Those positions were designed by FuncLib (*31*), using mutations with PSSM > −2 and ΔΔ*G* < 6 R.e.u. but present in the natural diversity of xylanases. The designs were computationally validated using AlphaFold2 (see above AlphaFold2 analysis of designed backbones) with 96.12 < plDDT < 98.13. The sequences can be found at <github>/data/small_set_xylanase.fa

#### Experimental validation of a small set of xylanases

The genes of the 27 designed xylanases were synthesized by TWIST Bioscience and cloned into pETMBPH vector using restriction sites *EcoRI* and *PstI*. The genes were transformed into BL21 DE3 cells and expressed in 10ml culture as described before. After centrifugation and storage at –20°C, the pellets were resuspended in the lysis buffer and lysed by sonication. Xylanase activity was assayed using 100 μl lysate and 0.5mM O-PNPX_2_ and cellPNP substrates, as described above, and seven most active designs were taken for further characterization (Table S4).

The proteins were expressed in 50 ml culture, lysed by sonication and the cleared lysate was bound to amylose resin (NEB), washed with 50mM Tris pH 7 with 100mM NaCl, and the proteins were eluted with wash buffer containing 10mM maltose. Elution fraction was analyzed for purity by SDS-PAGE and used for kinetic measurements. For stability measurements, the MBP fusion tag was cleaved by TEV (1:20 TEV, Tris 50mM with 100mM NaCl and 1mM DTT, 24-48h at room temperature), the MBP fusion tag was removed by binding to Ni-Nta resin.

#### Specific activity

(μM product per minute per mg protein) was measured with 0.5mM O-PNPX_2_ and cellPNP at pH 6.5 and 37°C by monitoring the absorbance of the leaving group at 405 nm. For determination of kinetic parameters, activity with O-PNPX_2_ was measured with a range of substrate concentrations, and the kinetic parameters were obtained by fitting the data to the Michaelis-Menten equation [v_0_ = *k_cat_*[E]_0_[S]_0_/([S]_0_ + *K*_M_)] using Prism 7. In cases where solubility limited substrate concentrations, data were fitted to the linear regime of the Michaelis-Menten model (v_0_= [S]_0_[E]_0_*k_cat_*/*K*_M_) and *k_cat_/K*_M_ values were deduced from the slope. The reported values represent the means ± S.D. of at least two independent measurements.

#### Activity with natural xylans

Xylanase activity with beechwood xylan and wheat arabinoxylan was determined by measuring the reducing sugars released from xylan by the dinitrosalicylic acid (DNS) method(*69*). 50μl protein was mixed with 50μl of 0.1M sodium citrate buffer pH 6, and added to 100μl of 2% xylan dispersed in water. Reaction mixture was incubated 0.5 hour at 50°C and then quenched for 10 minutes on ice. Then, 200μl of DNS reagent was added, the mixture was vortexed, incubated for 10 minutes at 95°C and cooled on ice. The mixture was diluted 1 to 4 in water, and absorbance at 540 nm was measured, compared to a blank sample (cell lysate expressing MBP). The reported values represent the means ± S.D. of two independent measurements.

#### T_M_ measurements

Melting temperatures of the tagless xylanases were measured by nanoDSF (nanoscale differential scanning fluorimetry) on Prometheus NT.Plex instrument (NanoTemper Technologies)(*70*). The temperature was increased from 20°C to 95°C at 1°C/min ramp, and the melting temperatures were calculated at inflection point.

### Supplementary Figures

**Fig. S1.**
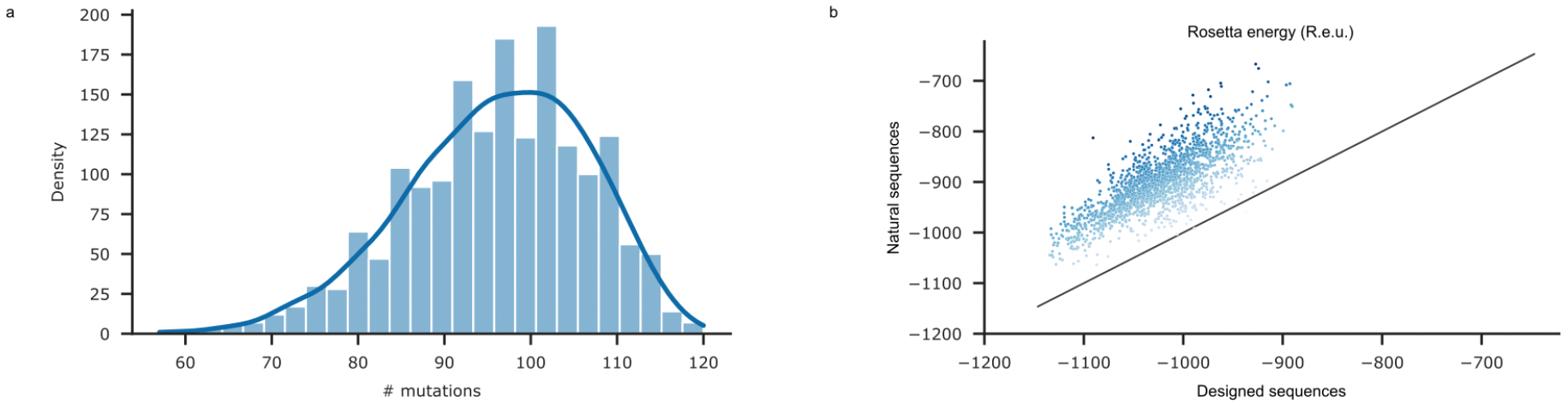
Designs carry a large number of stabilizing mutations. **(A)** Number of mutations in the set of designs assembled by selected fragments, compared to assemblies using the natural sequence of the fragments. **(B)** Comparing the Rosetta energies of designs assembled with designed fragments or natural sequences. Colors are darker shades in designs for which the energy difference is greater.

**Fig. S2.**
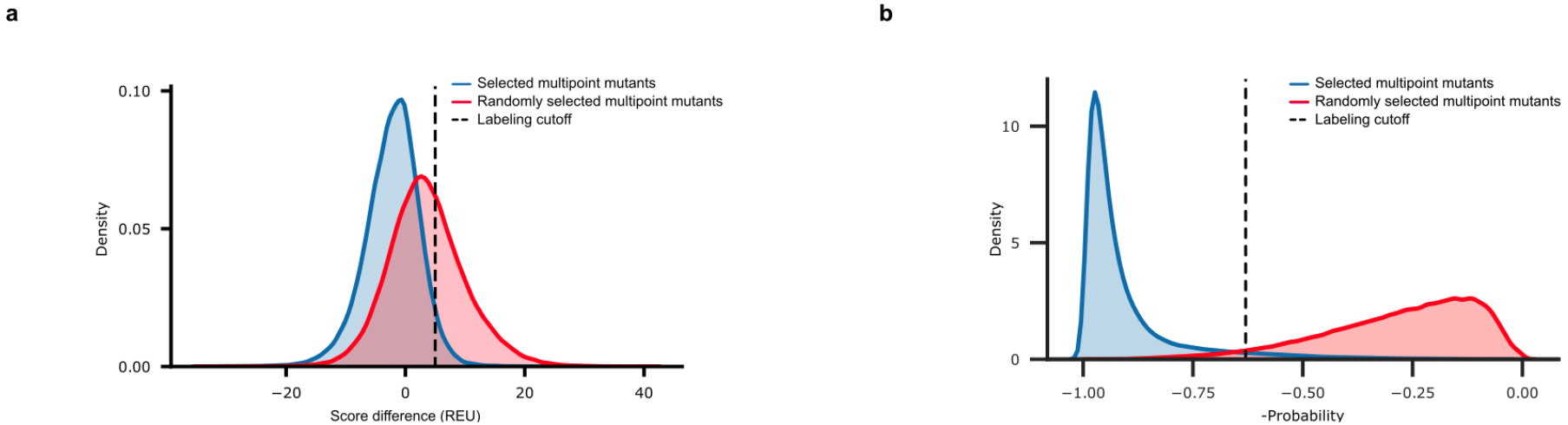
EpiNNet-based selection enriches for low-energy multipoint mutants in the sequence diversification step. **(A)** 100,000 randomly selected designs were modeled from either the multipoint mutants selected for the first library (blue) or from all multipoint mutants suggested by the PSSM and single-mutation energy (ΔΔ*G*) calculations (red). Multipoint mutants are labeled positive if their energy is up to 5 R.e.u. from the non-mutated design (dashed line). 96% and 62% designs are labeled positive in the selected and random populations, respectively. **(B)** 50,000 randomly selected designs were modeled from either the multipoint mutants selected for the second-generation library (blue) or from all multipoint mutants suggested by the ΔΔ*G* and PSSM cutoffs (red). Multipoint mutants are labeled positive if -probability < −0.63 (dashed black line). 96% and 4% of the designs are labeled positive in the selected and random populations, respectively.

**Fig. S3.**
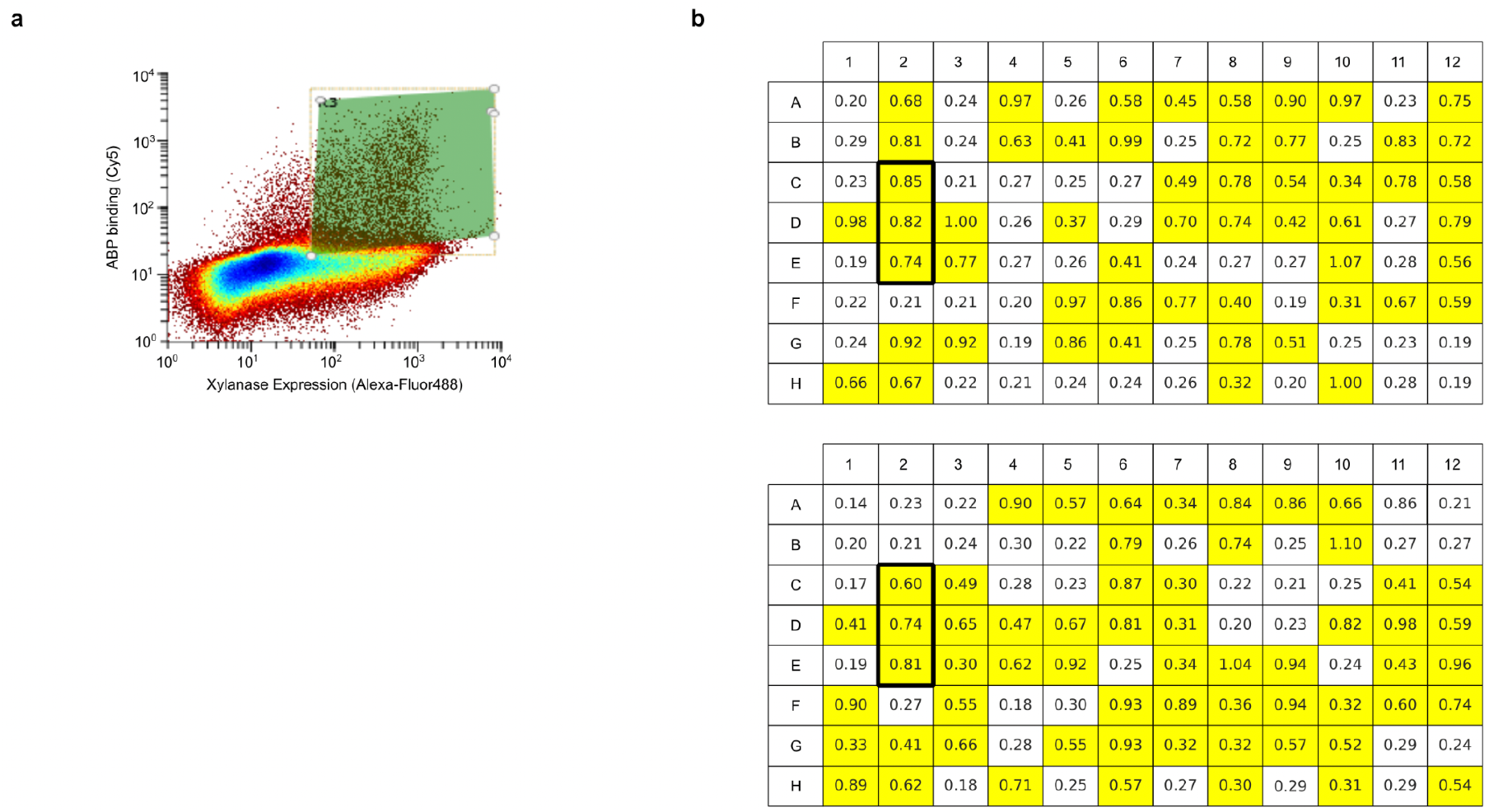
Design activity in the first-generation library. **(A)** Final FACS sort (sort 3) of the first-generation library with xyl-Cy5. The selection gate is shown in green. **(B)** OD_405_ of the individual clone lysates in 96-well plates, measured after overnight incubation with 0.5 mM OPNPX_2_. 58% of the variants were active (OD_405_>0.3). The boxed wells contained positive controls (see Methods).

**Fig. S4.**
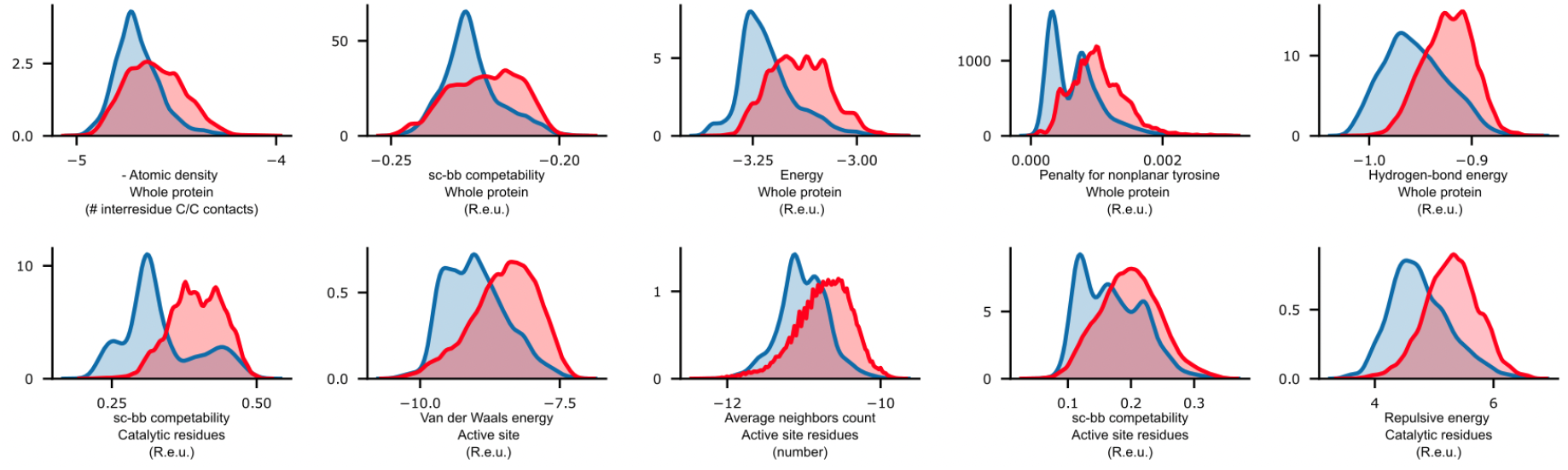
Features discriminating presumed active from inactive designs. The distribution of the features included in the activity predictors in the active and inactive design populations (blue and red, respectively). All features (except average neighbor count) are normalized by the number of residues.

**Fig. S5.**
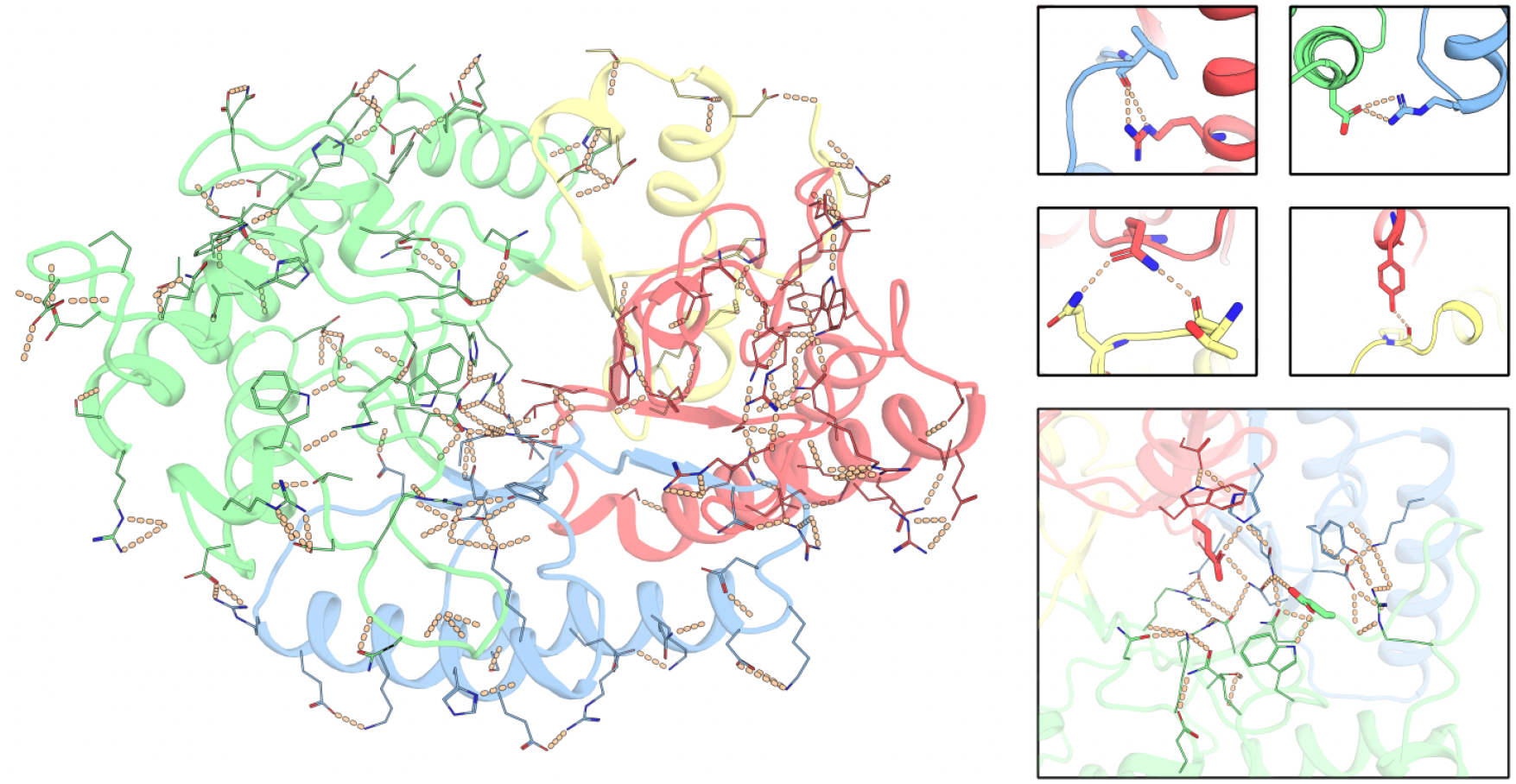
Hydrogen-bond networks in a representative designed TIM-barrel xylanase. **Left:** Showing hydrogen bonds involving sidechains in a designed active backbone (fragments sourced from PDB entries 4PUE, 3W23, 4PMD and 3MUI). **Right, top:** zoom-in to some hydrogen bonds involving residues from different segments. **Right, bottom:** zoom-in on hydrogen bond networks in the active site. Catalytic Glu residues in sticks.

**Fig. S6.**
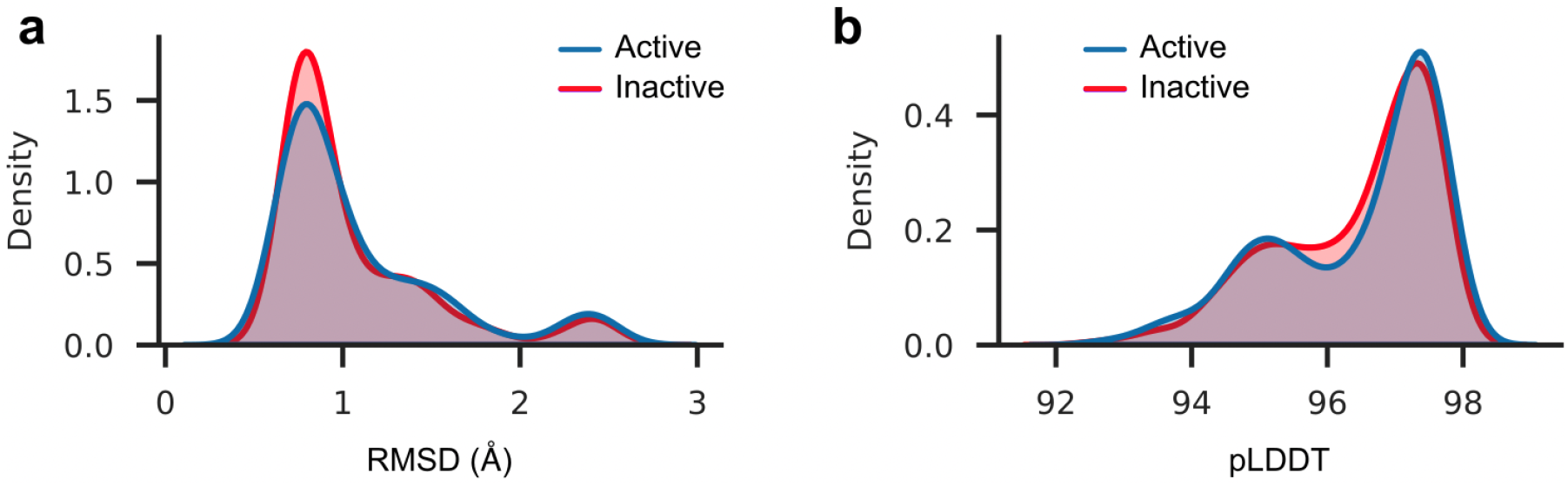
AlphaFold2 does not discriminate presumed active from inactive designs. **(A)** RMSD between Rosetta model structures and AlphaFold2 predicted structure for designs. Both active (blue) and inactive (red) backbones show similar distributions. **(B)** pLDDT values of predicted AlphaFold2 structures. Active and inactive backbones have similarly high average pLDDT scores.

**Fig. S7.**
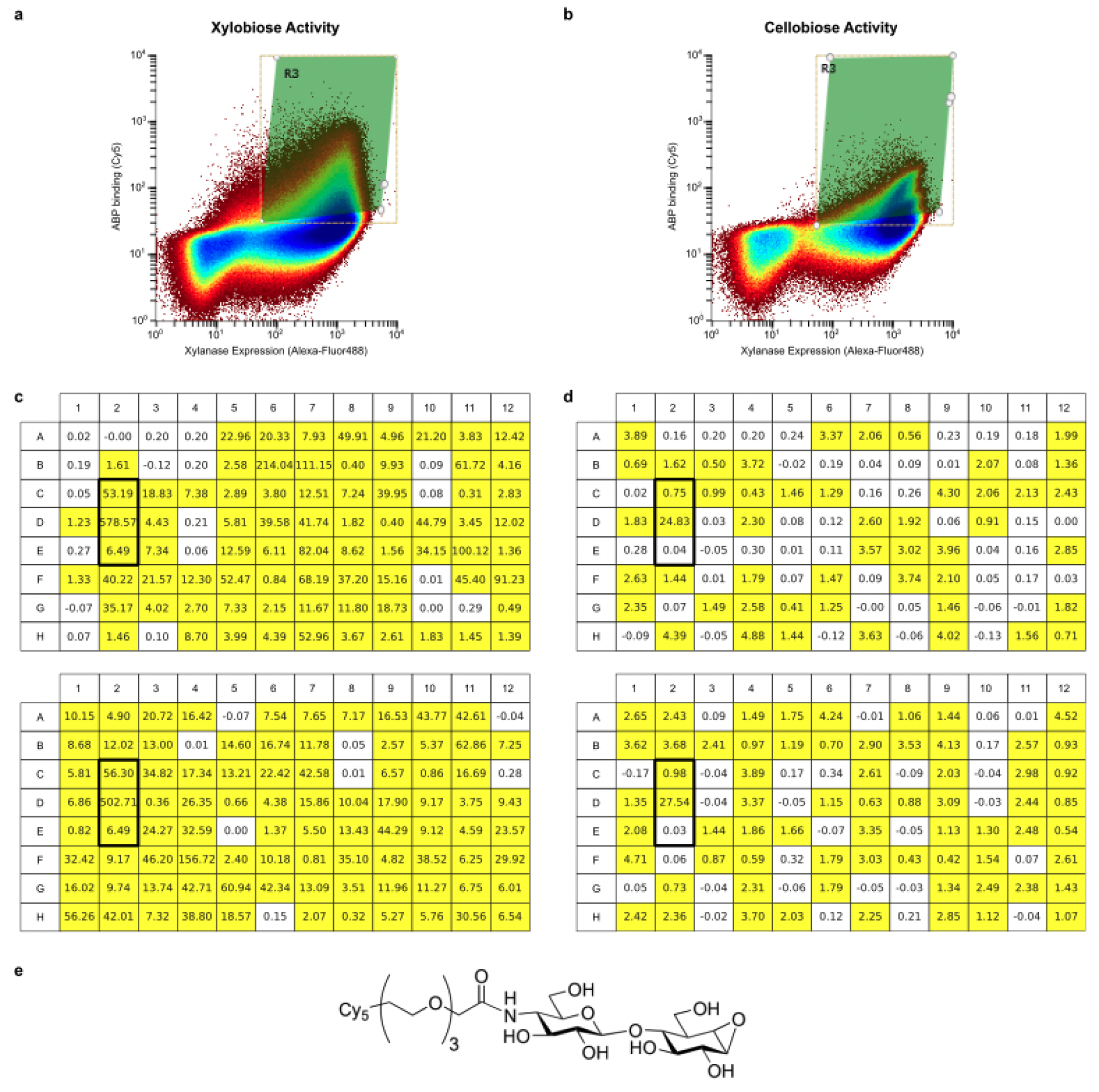
Catalytic activity in the second design round. **(A-B)** Final FACS sorts of the second-generation library. (a) Sort 3 with xyl-Cy5 and (b) sort 4 with cel-Cy5. **(C-D)** Initial reaction velocity (mOD/min) of the individual clone lysates in 96-well plates. **(C)** 85% of the variants were active with 0.5mM OPNPX_2_ (mOD/min > 0.3). **(D)** 60% of the variants were active with 0.5mM cellPNP (mOD/min > 0.4). The boxed wells contained positive controls (see Methods). **(E)** Structure of the cellobiose ABP.

### Supplementary Tables

**Table S1.**
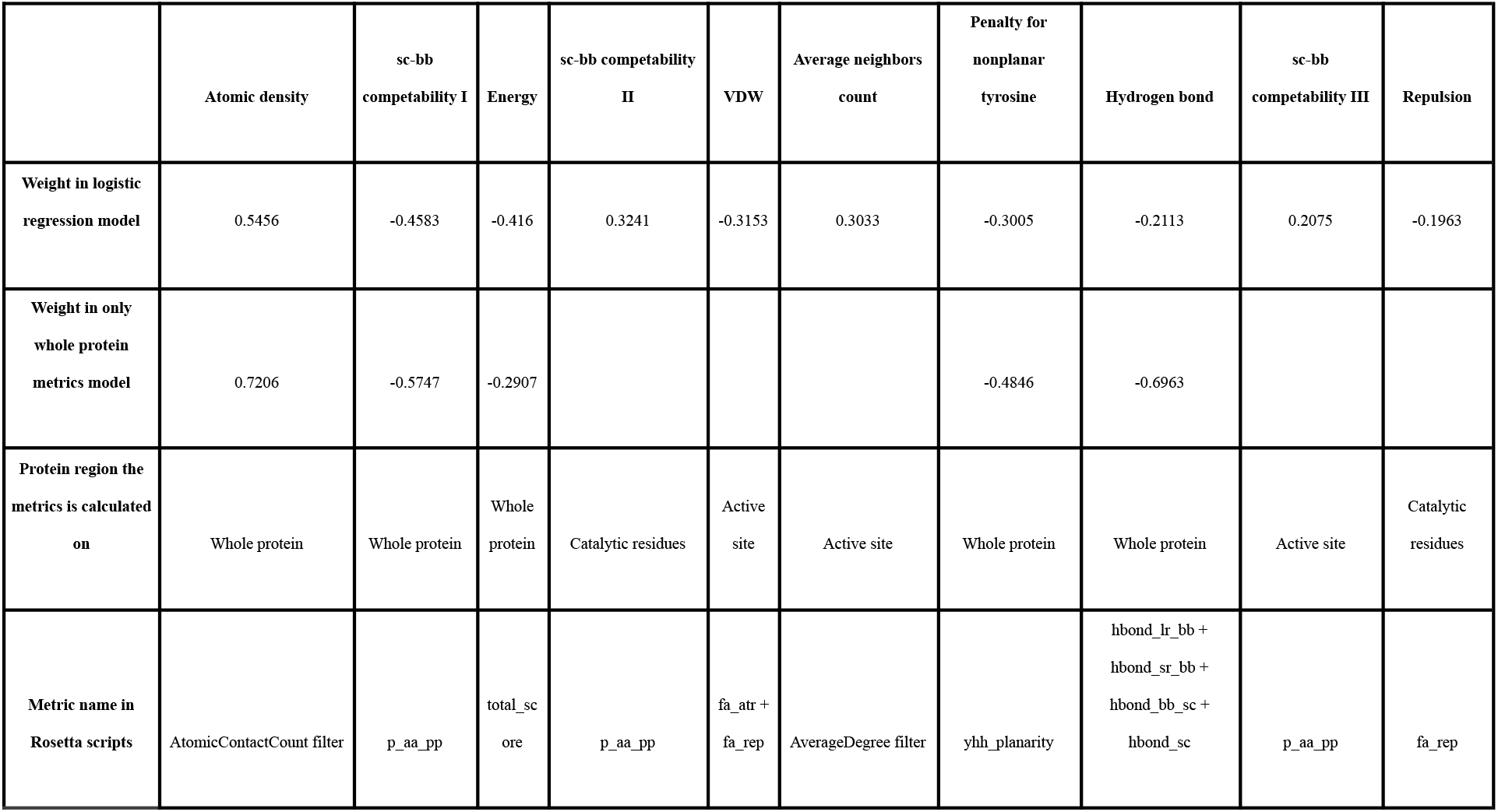
Features included in logistic regression models.

**Table S2.** Classification accuracy report on an independent test set for the activity predictor including active-site features.

|  | Precision | Recall | f1-score | support |
| --- | --- | --- | --- | --- |
| Inactive | 1 | 1 | 1 | 189792 |
| Active | 0.26 | 0.45 | 0.33 | 608 |
| accuracy |  |  | 0.99 | 190400 |
| macro avg | 0.63 | 0.72 | 0.66 | 190400 |
| weighted avg | 1 | 0.99 | 0.99 | 190400 |

**Table S3.** Classification report of the test set for activity predictor including only whole-protein features.

|  | Precision | Recall | f1-score | support |
| --- | --- | --- | --- | --- |
| Inactive | 1 | 0.99 | 1 | 189792 |
| Active | 0.19 | 0.43 | 0.27 | 608 |
| accuracy |  |  | 0.99 | 190400 |
| macro avg | 0.6 | 0.71 | 0.63 | 190400 |
| weighted avg | 1 | 0.99 | 0.99 | 190400 |

**Table S4.**
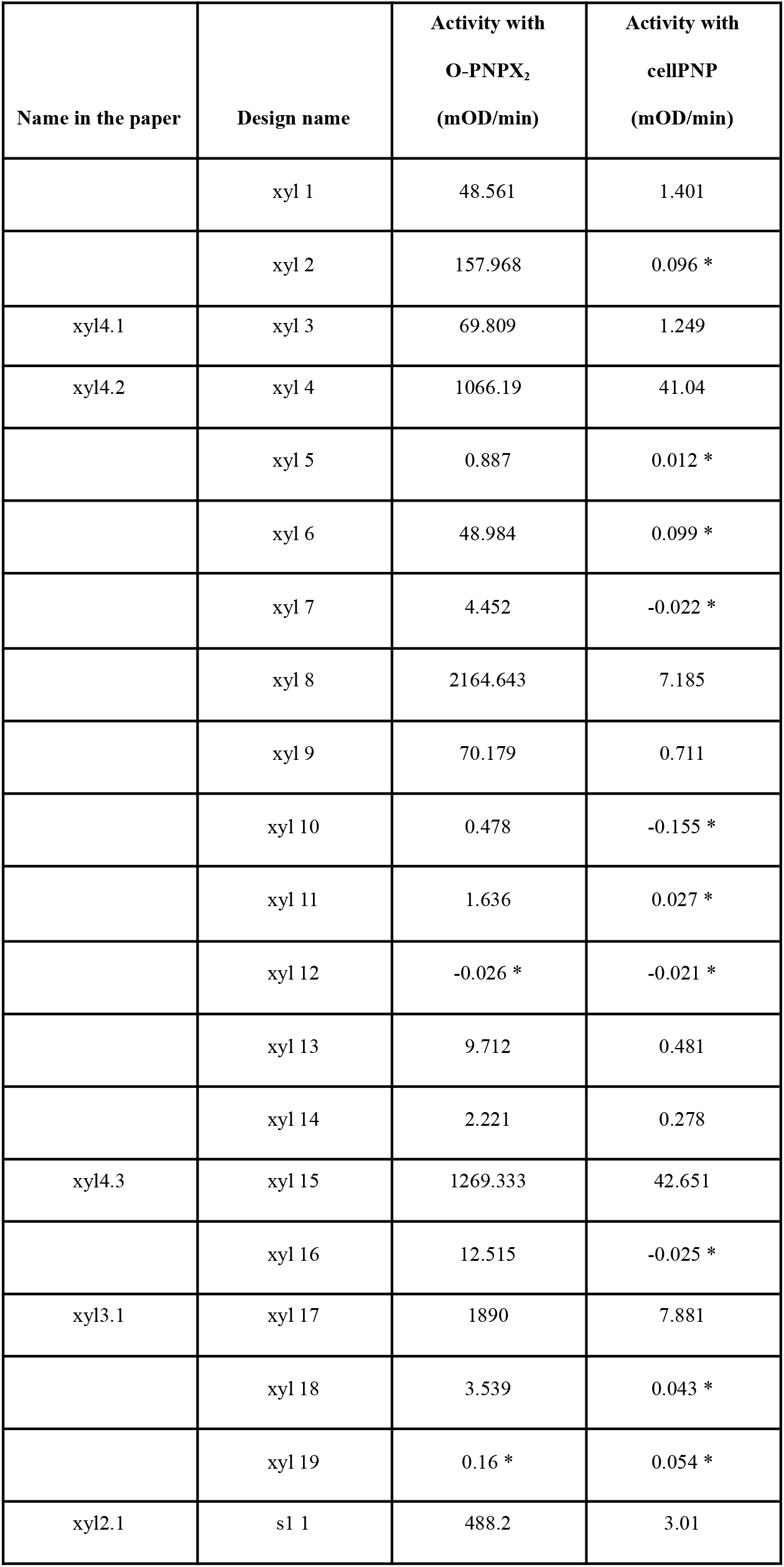
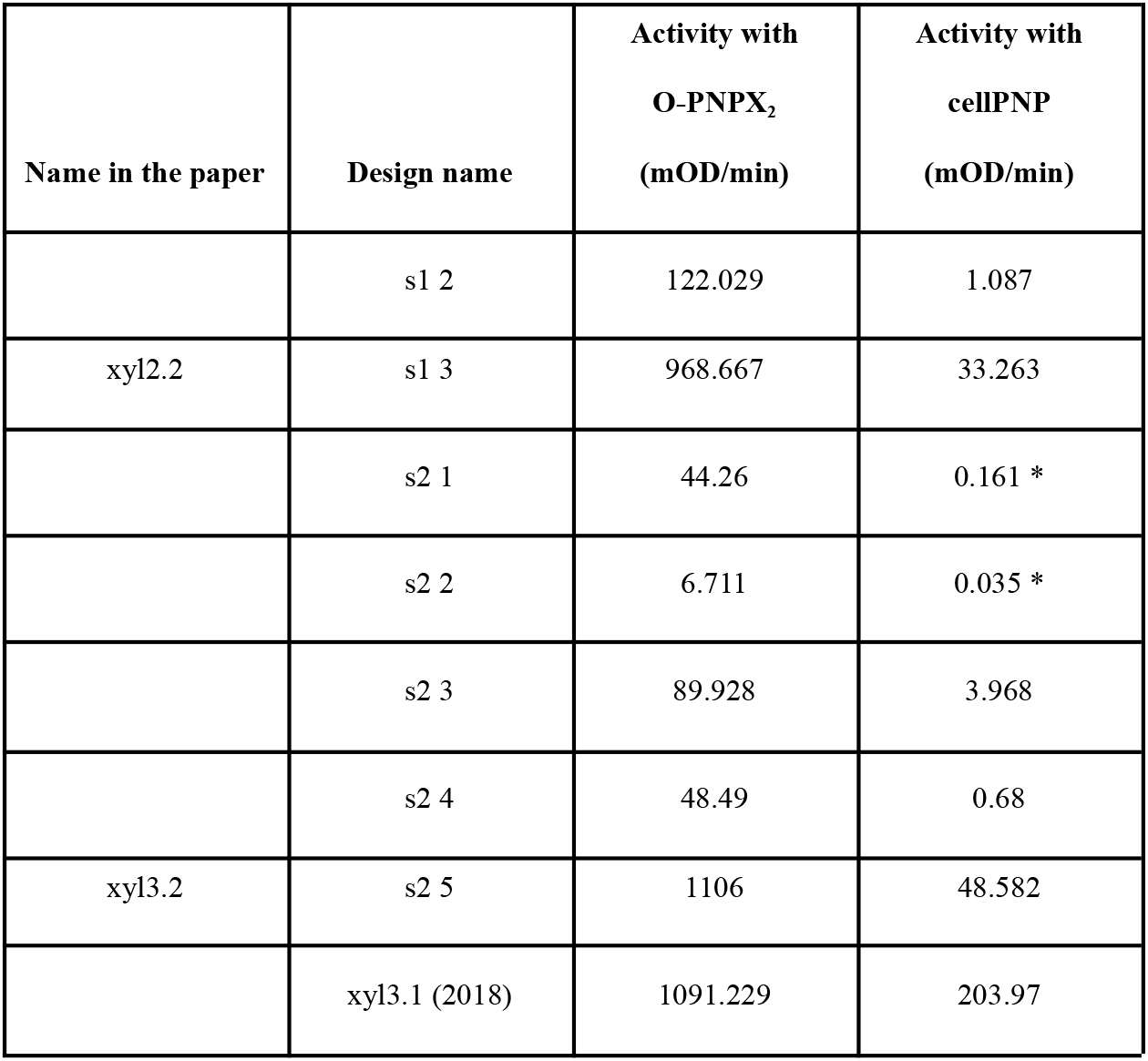
Initial reaction velocity of individual xylanase designs. Values marked with * showed no activity. Last row is the control (design xyl3.1 from(*18*))

| Name in the paper | Design name | Activity with<br>O-PNPX<sub>2</sub><br>(mOD/min) | Activity with<br>cellPNP<br>(mOD/min) |
| --- | --- | --- | --- |
|  | xyl 1 | 48.561 | 1.401 |
|  | xyl 2 | 157.968 | 0.096 * |
| xyl4.1 | xyl 3 | 69.809 | 1.249 |
| xyl4.2 | xyl 4 | 1066.19 | 41.04 |
|  | xyl 5 | 0.887 | 0.012 * |
|  | xyl 6 | 48.984 | 0.099 * |
|  | xyl 7 | 4.452 | -0.022 * |
|  | xyl 8 | 2164.643 | 7.185 |
|  | xyl 9 | 70.179 | 0.711 |
|  | xyl 10 | 0.478 | -0.155 * |
|  | xyl 11 | 1.636 | 0.027 * |
|  | xyl 12 | -0.026 * | -0.021 * |
|  | xyl 13 | 9.712 | 0.481 |
|  | xyl 14 | 2.221 | 0.278 |
| xyl4.3 | xyl 15 | 1269.333 | 42.651 |
|  | xyl 16 | 12.515 | -0.025 * |
| xyl3.1 | xyl 17 | 1890 | 7.881 |
|  | xyl 18 | 3.539 | 0.043 * |
|  | xyl 19 | 0.16 * | 0.054 * |
| xyl2.1 | s1 1 | 488.2 | 3.01 |
|  | s1 2 | 122.029 | 1.087 |
| xyl2.2 | s1 3 | 968.667 | 33.263 |
|  | s2 1 | 44.26 | 0.161 * |
|  | s2 2 | 6.711 | 0.035 * |
|  | s2 3 | 89.928 | 3.968 |
|  | s2 4 | 48.49 | 0.68 |
| xyl3.2 | s2 5 | 1106 | 48.582 |
|  | xyl3.1 (2018) | 1091.229 | 203.97 |

**Table S5.** Characterization of the most active individual xylanase designs: expression levels, stability and specific activity.

| **Design** | **Expression**<br>**mg protein/L culture** | **Stability**<br>**TM, °C (nanoDSF)** | **#aa difference from**<br>**nearest nr homolog** | **Specific activity**<br>**ratio O-PNPX<sub>2</sub> to cellPNP** |
| --- | --- | --- | --- | --- |
| xyl4.1 | 93.6 | 61.8 | 128 | 57 |
| xyl4.2 | 148.8 | 61.7 | 119 | 50 |
| xyl4.3 | 159 | 61.4 | 94 | 306 |
| xyl3.1 | 134.2 | 68.1 | 83 | 281 |
| xyl2.1 | 129.3 | 66.1 | 104 | 215 |
| xyl2.2 | 177 | 72.9 | 78 | 436 |
| xyl3.2 | 182 | 62.7 | 121 | 127 |
| 3W24 | 202.8 | 79.8 | - | 426 |

## Notes

https://github.com/Fleishman-Lab/CADENZ

### Competing Interest Statement

The authors have declared no competing interest.

### Summary of Updates

Title, abstract and discussion are revised

